# Tobacco Images Choice and Its Association with Craving and Dependence in Cigarette Smokers

**DOI:** 10.1101/2023.08.30.555462

**Authors:** Marcello Solinas, Claudia Chauvet, Claire Lafay-Chebassier, Paul Vanderkam, Lila Barillot, Scott J. Moeller, Rita Z. Goldstein, Xavier Noël, Nematollah Jaafari, Armand Chatard

**Affiliations:** Laboratoire de Neurosciences Expérimentales et Cliniques, Université de Poitiers, INSERM, U-1084, Poitiers, France; Unité de Recherche Clinique Pierre Deniker, Centre Hospitalier Henri Laborit, Poitiers; INSERM, Centre d’Investigation Clinique CIC 1402, Université de Poitiers, CHU Poitiers, Poitiers, France; Service de Pharmacologie Clinique, CHU Poitiers, Poitiers, France; Université de Poitiers, Université de Tours, CNRS 7295, CeRCA, Poitiers, France. France; Department of Psychiatry and Behavioral Health, Renaissance School of Medicine at Stony Brook University, Stony Brook, NY 11794, USA; Psychiatry & Neuroscience, Icahn School of Medicine at Mount Sinai, NY, USA; Université Libres de Bruxelles, Belgium; Service Hospitalo-Universitaire de Psychiatrie et de Psychologie Médicale, Centre Hospitalier Henri Laborit, Poitiers, France

## Abstract

**Introduction:** Increased salience of drug-related cues over non-drug reinforcers can drive drug use and contribute to tobacco use disorder (TUD). An important scientific and clinical goal is to effectively measure this elevated drug-seeking behavior in TUD. However, most TUD assessments rely on self-reported cravings and cigarette consumption, not providing an objective measure of the impact of drug-cues on biasing behavior towards drugs. The probabilistic image choice (PIC) task investigates the choice of viewing drug-related pictures as compared to other salient pictures (e.g., pleasant and unpleasant). This study aimed to develop and validate the PIC task for TUD and evaluate the associations between behavioral choice and tobacco craving, daily cigarette consumption, quit attempts and motivation to quit, and nicotine dependence (the Fagerström score).

**Methods:** We recruited 468 smokers and 121 nonsmokers using the Prolific online platform. Participants performed the PIC task twice (at a one-month interval) and completed other measures relevant to TUD.

**Results:** compared to nonsmokers, tobacco smokers selected to view significantly more tobacco images and less pleasant (non-drug reinforcer) images, a profile that remained stable at retest. Individual differences in choice of tobacco as compared to pleasant images on the PIC task were associated with craving but not with the other tobacco dependence measures, suggesting that the task may serve as a behavioral proxy measure of drug “wanting” rather than of cumulative nicotine exposure or physical dependence.

**Conclusions:** these results suggest that the PIC task can be a valuable tool for objectively assessing craving-associated tobacco seeking in TUD.

**Implications section**, which should provide a brief description about what the study adds.

Most of the current measures of tobacco use disorder (TUD) rely on self-reports of consumption, dependence and craving and do not take into consideration the role of drug-related cues in driving tobacco seeking. This study shows that the probabilistic image choice (PIC) task provides an objective, reliable proxy measure of tobacco image seeking behavior in cigarette smokers that is linked to craving (desire) for smoking but not to other measures of TUD. Therefore, the PIC task may be a useful complementary tool for the classification, diagnosis, and prognosis of TUD.

## INTRODUCTION

Tobacco use disorder (TUD) is the major preventable cause of morbidity and mortality in the world ^1^. In the last decades, prevention campaigns have been effective in educating the general population about the risks associated with tobacco smoking, contributing to reduced prevalence of TUD ^2,3^. However, with the increase in global population, the total number of smokers has increased and TUD is estimated to cause 7 million excess deaths per year ^2,3^. Investigating the factors that drive tobacco use can provide a deeper understanding of the mechanisms underlying TUD and help health-care providers to improve intervention and prevention efforts for this treatment-resistant disorder.

A critical aspect of TUD is relapse, occurring within a year of a quit attempt in more than 50% of abstinent smokers ^4^. Drug-related cues play a major role in relapse and the maintenance of drug addiction ^5–10^. Indeed, a contributing factor are the strong associations formed between the drugs’ reinforcing effects and environmental cues associated with drug use ^11,12^. This process underlies the attribution of excessive value to these cues that become irresistible “motivational magnets” driving drug seeking behavior ^13^ even when awareness of such choice is compromised ^14–16^. Moreover, over time the value of drugs and related cues increases at the expense of alternative reinforcers ^17^. Drug-biased incentive motivation is also important for TUD because, although nicotine is a relative weak reinforcer in animal models of addiction, it increases the incentive properties of nicotine-associated cues so that they can act as reinforcers even in the absence of nicotine ^18^. Nevertheless, aside from tobacco self-administration paradigms ^19^, impractical and/or unethical in many settings, current measures of TUD such as the Fagerström Test for Nicotine Dependence (FTND) rely mostly on self-reported measures of craving, drug use and dependence and do not provide an objective measure of cue-induced drug seeking. The use of operant measures of the impact of drug-cues on biasing behavior towards drugs can help ascertain the severity of addiction ^20^, enhancing the translational nature of such human research. This approach also aligns with the recommendation to incorporate objectively-measured outcome variables in clinical trials focusing on drug addiction^21^.

Within the framework of behavioral economic theories of addiction ^22,23^, choice procedures using binary and mutually exclusive choices have been developed to investigate the extent and nature of drug users’ preferences for drugs over alternative rewards, identifying the factors that influence these choices ^22,24–26^. However, in real life conditions, people are rarely confronted with such choices; instead, they usually face multiple options associated with probabilistic outcomes. This creates a context where people may not always be fully aware of their own choices, which is particularly important in as much as compromised self-awareness is thought to play a major role in addiction ^14,16,27–30^. Therefore, developing procedures that allow measuring cue-induced drug-related choice behavior in a probabilistic context could be helpful in identifying individuals who can benefit from targeted behavioral interventions (e.g., training to enhance self-awareness of choice, developing strategies to effectively regain control over behavior).

The probabilistic image choice (PIC) task was originally designed as a tool to objectively model drug-related choice behavior in individuals with cocaine use disorder (both active and abstinent users) ^31^. Participants make selections between flipped-over cards included in four decks where image identity (pleasant, unpleasant, neutral and drug) is both not certain and varies as the task progresses, allowing tapping into more implicit processes linked to choice. Results of previous studies using this task showed higher preference for viewing cocaine pictures in people with cocaine use disorder than healthy controls, driven by current users but observable, to a lesser extent, also in those abstaining from cocaine ^31^. Importantly, the selection of cocaine over pleasant images (the alternative positive reinforcer) was associated with both current and predictive of future drug use, pointing to a direct comparison that is crucial for identifying individual differences in choice relevant to actual drug seeking ^32^. Although the PIC task has been successfully adapted to methamphetamine ^33^ and opiate ^34^ use disorders, its potential use for investigating TUD has not been explored.

In this study, we adapted the PIC task to tobacco smoking and investigated differences between 468 smokers and 121 non-smokers, as well as test-retest stability of tobacco image seeking behavior after a one-month interval. We further explored whether tobacco-seeking behavior on the PIC task correlated with other measures of TUD such as craving, cigarette consumption, quit attempts and motivation to quit, and severity of dependence. We hypothesized that, compared to non-smokers, smokers would choose more tobacco-related images, and less pleasant images (indicative of a drug>pleasant shift in valuation and behavior, and consistent with our prior reports in other addictions). Additionally, we hypothesized that a preference for tobacco images would be associated with self-reported measures of craving, daily cigarette consumption, quit attempt/motivation to quit, and severity of tobacco dependence.

## METHODS

### Participants

Participants were 468 regular smokers (i.e., individuals who smoked at least 5 cigarettes per day over the last year) and 121 nonsmokers (i.e., never smokers) recruited via the research online platform Prolific (https://www.prolific.com/), see Table 1 for sample demographics). Participants were eligible for the study if they were residents from the UK or the USA, and if they were native English speakers. When included in the analysis, the country did not moderate any effects reported in the text. Therefore data from USA and UK participants were pooled. In an effort to oversample individuals who smoked cigarettes for several years, only participants older than 35 were recruited. Smokers completed the study on March 3^rd^ 2022 (Time 1). About 30 days later, on April 4^th^ 2022, the participants were recontacted via the platform Prolific and invited to respond to a follow-up survey (Time 2). A total of 286 participants completed the second wave of data collection (182 participants could not be reached, were not available, or did not respond to the survey). Of these, 13 participants were excluded because they reported smoking less than 5 cigarettes. Nonsmokers completed the study on February 2^nd^, 2023. All participants were debriefed and received 10£/hour as compensation.

**Table 1.** Demographic characteristics.

| Smoking Status | Age ± SD (Min-Max) | Gender | Frequency | Percent | Valid Percent |
| --- | --- | --- | --- | --- | --- |
| Smokers | 47.47 ± 9.50 (35-75) | Female | 206 | 44.02 | 49.16 |
|  |  | Male | 211 | 45.08 | 50.36 |
|  |  | Nonbinary | 2 | 0.43 | 0.48 |
|  |  | Missing | 49 | 10.47 |  |
|  |  | Total | 468 | 100.00 |  |
| Nonsmokers | 50.58 ± 10.46 (34-76) | Female | 60 | 49.59 | 50.42 |
|  |  | Male | 59 | 48.76 | 49.58 |
|  |  | Nonbinary | 0 | 0.00 | 0.00 |
|  |  | Missing | 2 | 1.65 |  |
|  |  | Total | 121 | 100.00 |  |

The study was approved by the local institutional ethical committee (Comité d’Ethique de la Recherche des Universités de Tours et Poitiers) and conforms with the European regulations on data protection.

### Procedure

Participants were initially classified as smokers or nonsmokers based on information in the Prolific database. After a few demographic questions (gender, age, nationality, etc.), smokers provided information about their tobacco consumption (the number of cigarettes smoked per day), quit attempts and motivation to quit followed by a self-reported questionnaire on craving^35^. Then all participants completed the PIC task. Finally, smokers were asked to complete the Fagerström test for nicotine dependence ^36^. These measures are described in detail below. Our study incorporated several attention checks to ensure that participants were actively engaged and followed the instructions accurately. The vast majority (99%) of participants successfully cleared these checks. This result aligns with recent studies indicating that Prolific participants tend to perform tasks at a high quality level ^37^. Participants who did not pass were preemptively excluded through Prolific’s filtering process. That is, all participants included in the dataset had successfully passed the attention checks.

### Probabilistic Image Choice (PIC) Task

The ‘probabilistic’ version of the original choice task developed by Moeller et al. (2009) was adapted and used in the present study to measure the behavioral tendency to choose drug as compared to other salient (pleasant and unpleasant) and neutral pictures. Participants were presented with four decks of flipped-over cards. Participants were informed that each deck contains neutral, pleasant, unpleasant, and drug-related pictures, but that some decks contain more pictures of one type than the others. Participants were asked to find and select the deck(s) that they found most appealing. Upon selection, the picture was uncovered to fit the entire screen and passively viewed for 2000 ms (Fig. S1). Participants could then select again from the same deck or switch to another one. Each deck contained a total of 30 pictures, which unbeknownst to the participants were pseudorandomly sorted according to the following two constraints: (A) there were no picture repetitions between the four decks; and (B) each deck contained 26 pictures (87%) of one picture category (e.g., drug: tobacco), two pictures (7%) of another category (e.g., pleasant), and one picture (3%) of each of the remaining two categories (e.g., unpleasant, neutral). These percentages were selected to reduce awareness of deck identity, while still allowing for preference to be established. A run terminated when participants selected from a particular deck for a total of eight times. Participants completed four runs. To further reduce awareness of deck identity, and to overcome the potential impact on results of perseverative responding (e.g., repeatedly choosing from the same deck across the runs), the dominant picture categories were pseudorandomized across the decks between the runs (i.e., the deck location of the four picture categories did not repeat across the runs). The total number of cards selected per picture category (neutral, pleasant, unpleasant, and tobacco-related) across the four runs was summed. As in previous studies ^31,32^, drug seeking was operationalized as the preference for viewing tobacco directly as compared to the pleasant images.

We replaced the pictures used in the previous PIC publications for improved picture quality and to fit online administration to people residing across two continents. The current version of the task, adapted for TUD, included 30 pleasant, 30 neutral, and 30 unpleasant pictures selected from the International Affective Picture System (Lang et al., 1998), depicting pleasant scenes (e.g., smiling faces, baby images, natural landscapes), neutral scenes (e.g., places, household objects), or unpleasant scenes (e.g., sad faces, violent images), respectively. The fourth picture category comprised of 30 images related to smoking cigarettes, tobacco, and tobacco use (e.g., cigarette packs, cigarette smoke, people smoking) derived from the Geneva Smoking Pictures library ^38^. The current version of the task was initially developed in Python and later converted to JavaScript to enable online functionality. The task is freely accessible via the Open Science Framework link: https://osf.io/n23rx/.

### Cigarettes consumption

Smokers were requested to report the quantity of cigarettes they typically smoke daily at both study sessions. Because this measure was heavily skewed to the right, it was transformed into a 4-point response scale, mirroring the one utilized in the Fagerström test (≤10 cigarettes per day = 0, 11 to 20 cigarettes per day = 1, 21 to 30 cigarettes per day = 2, and ≥31 cigarettes per day = 3). At Time 1, participants also indicated whether they have already tried to quit smoking in the past and to estimate the number of times they attempted to quit smoking. Finally, participants reported their motivation to quit smoking in the next 6 months on a 10-point Likert scale ranging from 1 = very weak to 10 = very strong.

### Craving questionnaire

The brief questionnaire of smoking urges (QSU-brief; ^35^) was used at both sessions to measure craving for cigarettes. This 10-item instrument assesses the desire to consume cigarettes in the immediate present (representative items: “All I want right now is a cigarette” and “I have an urge for a cigarette”). Participants indicated their response to each item using a 7-point Likert scale (1 = strongly disagree and 7 = strongly agree). In the present sample, the internal validity of the scale was satisfactory (Cronbach’s alpha = .95 and .96, at Time 1 and Time 2, respectively). The responses to the 10 items were summed to form a composite score of craving. We utilized the total score of the QSU-Brief due to its high internal consistency, suggesting a cohesive construct across all items. However, also we conducted an exploratory factor analysis with promax rotation, in line with Toll et al. (2006), to investigate the distinct contributions of the two subscales ^39^. This analysis validated the two-factor structure and revealed a strong correlation between these factors in our sample at T1 (r = .81, p < .001) and T2 (r = .85, p < .001). Subsequently, we replicated the analyses detailed in the manuscript for each subscale separately and found that the results were consistent across both factors and closely aligned with those obtained using the total score.

### Nicotine dependence

At Time 1, participants also completed the Fagerström Test for Nicotine Dependence ^36^. This is a standard instrument for assessing the severity of addiction to nicotine, providing an ordinal measure of nicotine dependence related to cigarette smoking. It contains six items that evaluate the quantity of cigarette consumption, the compulsion to use, and dependence (e.g., “Do you find it difficult to refrain from smoking in places where it is forbidden (e.g., in church, at the library, in the cinema?”). The six items were summed to yield a total score of 0-10.

### Statistical analyses

Prior to the main analyses, we screened for outliers using a ROUT test, setting the default value of Q (the threshold to determine the likelihood of being considered an outlier) at 1%; no outliers were detected. For the main analyses, at Time 1, we conducted a mixed 2 (Smoking status: smoker versus nonsmoker) x 4 (picture type: tobacco-related, pleasant, unpleasant, and neutral) analysis of variance (ANOVA) with repeated measures on the latter factor to examine whether TUD is associated with a higher preference for tobacco-related stimuli in the PIC task. We performed traditional analyses of variance (ANOVA) to align with previous research on the PIC task and to allow for direct comparisons with those studies. However, robust ANOVA, with a trimming level of 0.2, produced analogous results. To avoid redundancy and maintain consistency with prior studies, we decided to report only the ANOVA results in the text. We followed up significant interactions with post-hoc t-tests. Intraclass correlation (ICC) coefficients were also calculated to assess the stability of tobacco image seeking behavior and craving after a 1-month follow-up. Furthermore, we conducted Person’s correlations between the normalized and a priori designated metric indexing preference for viewing tobacco images over pleasant images (i.e., number of selections of tobacco images minus pleasant images) with all TUD-related measures (encompassing severity of cigarette consumption, craving, quit attempts and motivation to quit, and the Fagerström score). Because choice of tobacco>pleasant images correlated with the intensity of cravings (as detailed in the Results, below), we used a median split to create two sub-groups: individuals with high cravings and those with low cravings. Within the smokers, we then conducted a mixed 2 (Craving levels: High or Low) x 4 (picture types) ANOVA with repeated measures, separately for Time 1 and 2. This analysis was used to examine whether craving levels influenced image choice in the adapted PIC task. All data analyses were performed with R version 3.6.1 (R Core Team, 2021).

## RESULTS

Table 2 summarizes the main tobacco-related information in the population of smokers involved in our study.

**Table 2.** Descriptive Statistics of TUD-related measures.

|  | Number of<br>Cigarettes<br>T1 | Number of<br>Cigarettes<br>T2 | Number<br>of Quit<br>Attempts<br>T1 | Number of<br>Quit<br>Attempts<br>T2 | Quit<br>Motivation<br>T1 | Fagerström<br>T1 | Craving<br>T1 | Craving<br>T2 |
| --- | --- | --- | --- | --- | --- | --- | --- | --- |
| Valid N | 453 | 271 | 430 | 250 | 415 | 468 | 468 | 286 |
| Mean | 15.17 | 15.35 | 3.02 | 3.19 | 5.19 | 4.39 | 37.71 | 36.87 |
| SD | 7.64 | 7.57 | 3.21 | 2.98 | 2.81 | 1.95 | 15.27 | 16.79 |
| Minimum | 5.00 | 5.00 | 0.00 | 0.00 | 0.00 | 0.00 | 10.07 | 10.03 |
| Maximum | 50.00 | 50.00 | 25.00 | 15.00 | 10.00 | 10.00 | 69.92 | 69.85 |

At time 1 (baseline), the 2 (smoking status) x 4 (picture type) mixed ANOVA showed no group main effect of smoking status (*F*(1, 586) = 0.014, *p* = 0.90, η^2^_p_ = 0.001). However, there was a significant main effect of picture type (*F*(3, 1758) = 170.31, *p* < 0.0001, η^2^_p_ = 0.225), indicating that all participants chose more pleasant images than any of the other picture types (all pair-wise comparison tests, *p*s < .001). Of greater interest, this main effect was qualified as expected by a significant interaction (*F*(3, 1758) = 8.10, *p* < 0.001, η^2^_p_ = 0.014) (Fig. 1). Post-hoc tests revealed that while there were no significant differences between smokers and non-smokers in the selection of neutral (*p* = .144, Cohen’s *d* = -.15) or unpleasant images (*p* = .560, Cohen’s *d* = .05), tobacco smokers selected significantly more tobacco-related images (*p* < .001, Cohen’s *d* = .29) and significantly fewer pleasant images (*p* = .036, Cohen’s *d* = -.24) than non-smokers.

**Fig. 1.**
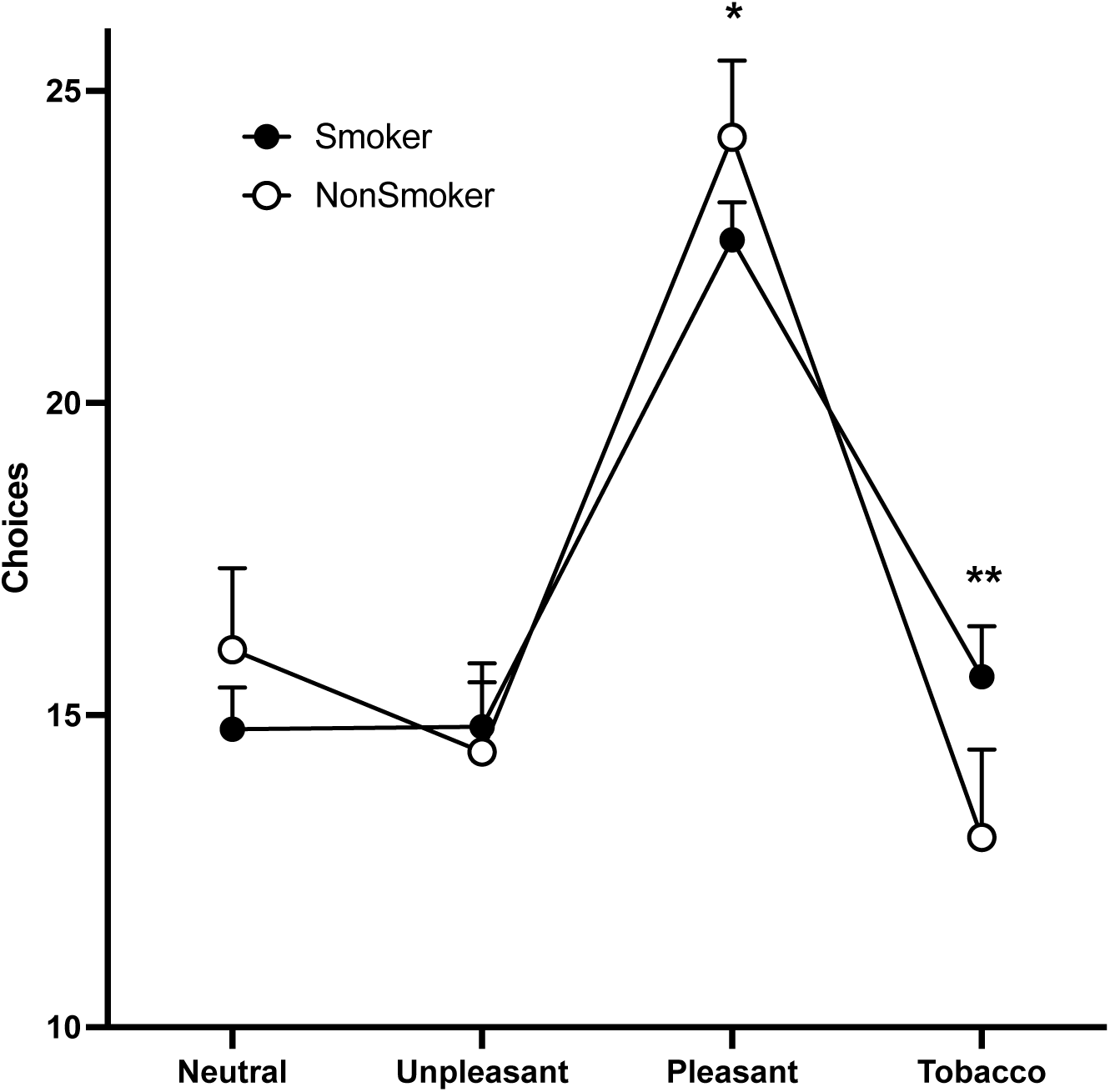
Performance in the PIC task in smokers and nonsmokers. A) Number of selection of images in each category in the PIC task for smokers (full circles, N = 468) and nonsmokers (empty circles, N = 121). Data are represented as mean ± 95% CI. * and **, *p* < 0.05 and *p* < 0.001 smokers vs. nonsmokers, respectively.

Within the smokers, the preference for viewing tobacco compared to other positively valenced images (number of selections of tobacco images minus pleasant images) was correlated across both Time 1 and Time 2 (ICC = .70, *p* < .0001; mean difference between timepoints: Cohens’ dz = .18), indicating that the probabilistic choice test shows sufficient test-rested reliability (Fig. 2). There were also high intercorrelations between T1 and T2 for craving scores (*r* (286) = .72, *p* < .001), number of cigarettes (*r* (286) = .88, *p* < .001) and quit attempts (*r* (286) = .81, *p* < .001).

**Fig. 2.**
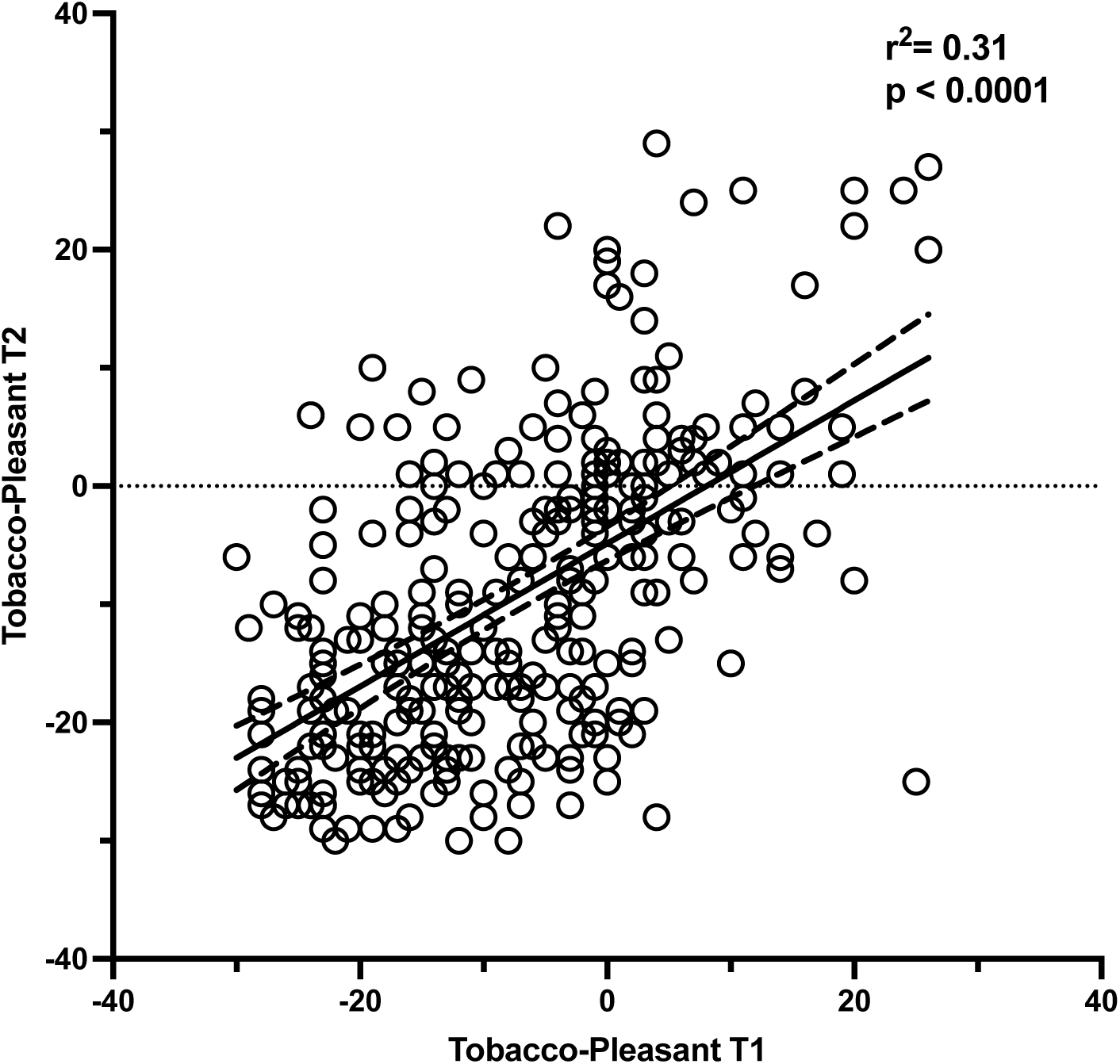
Test-retest reliability of tobacco choice on the PIC task. Correlation between the direct contrast of the selection of tobacco vs. pleasant images on the PIC task at Time 1 and Time 2, separated by a one-month interval, in smokers (N = 286). Dotted lines show 95% CI.

**Fig. 3.**
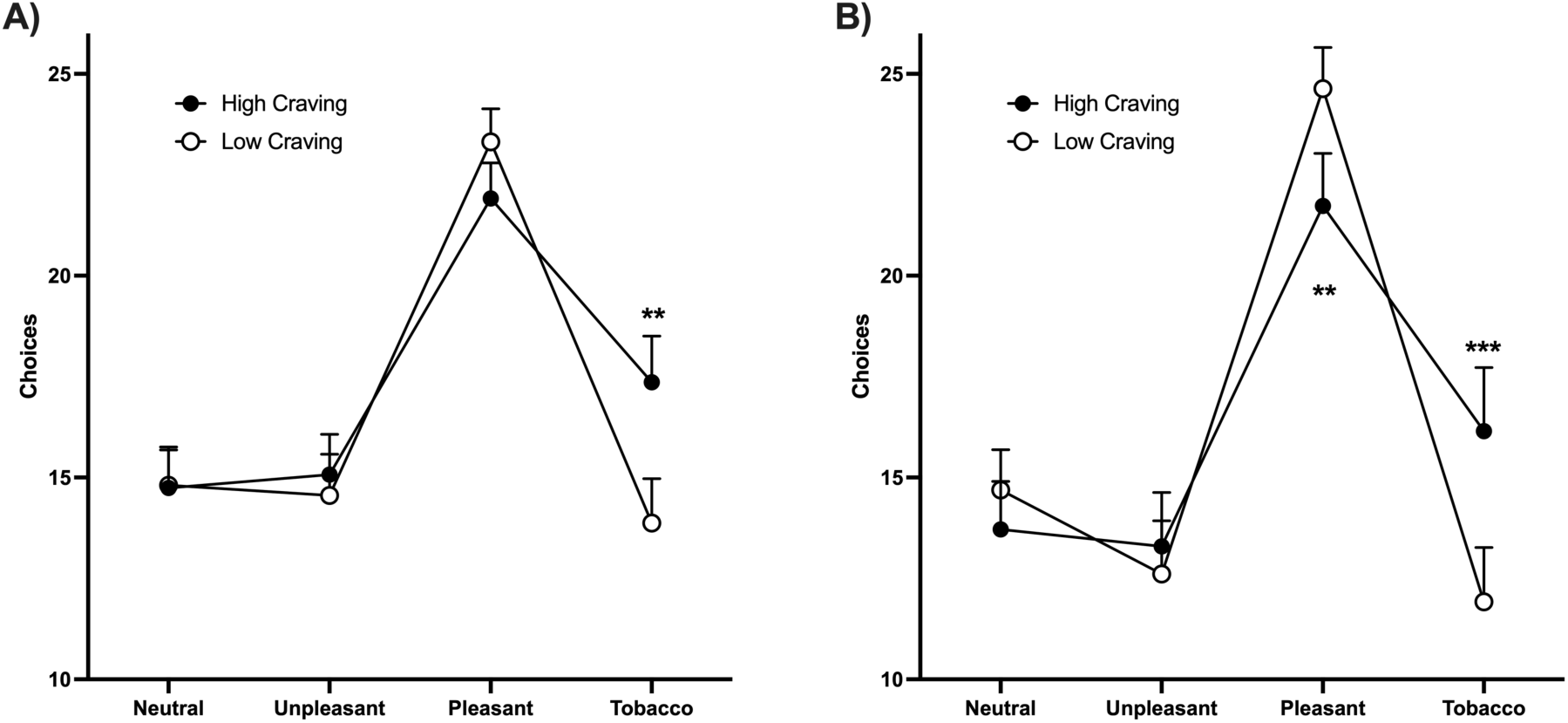
Performance in the PIC task in smokers as a function of craving. A) Number of selection of images in each category in the PIC task for high craving (full circles, N = 234) and low craving (empty circles, N = 234) smokers at T1. B) Number of selection of images in each category in the PIC task for high craving (full circles, N = 143) and low craving (empty circles, N = 143) smokers at T2. Data are represented as mean ± 95% CI. **, p < 0.001 high craving vs low craving.

Among the smokers, we also investigated Pearson correlations between our selected dependent measure (tobacco vs. pleasant viewing choice) with the tobacco-related variables (table 3). Tobacco>pleasant choices were positively correlated with craving both at time 1 (*r* (468) = .215, *p* < .001) and at time 2 (*r* (286) = .348, *p* < .001) (the higher the self-reported craving, the higher the tobacco>pleasant choice) (Fig. S2). In addition, drug>pleasant choices at time 1 prospectively predicted craving at time 2 (*r* (286) = .286, *p* < .001). Interestingly, this correlation remains significant even after partializing out the effect of craving at time 1, with rp (286) = .146, p = .014. Tobacco>pleasant choice was uncorrelated with the other TUD-related measures, at either T1 (r<0.098) or T2 (r<0.126). For correlations within the other tobacco use measures see Table 3.

**Table 3.** Pearson’s Correla0ons (and p-values) of TUD-related measures.

|  | T1–<br>Tobacco-Pleasant<br>Preference | T1–<br>Numbers of<br>Cigarettes | T1–Quit<br>Attempts | T1–Quit<br>Motivation | T1–<br>Fagerström | T1–Craving | T2–<br>Tobacco-Pleasant<br>Preference | T2–<br>Numbers of<br>Cigarettes | T2–Quit<br>Attempts | T2–Craving |
| --- | --- | --- | --- | --- | --- | --- | --- | --- | --- | --- |
| T1–Tobacco-Pleasant<br>Preference | — |  |  |  |  |  |  |  |  |  |
| T1–Numbers of<br>Cigarettes | 0.013 | — |  |  |  |  |  |  |  |  |
| T1–Quit Attempts | –0.060 | 0.017 | — |  |  |  |  |  |  |  |
| T1–Quit Motivation | –0.098<sup>+</sup> | **–0.104<sup>\*</sup>** | **0.356<sup>\*\*\*</sup>** | — |  |  |  |  |  |  |
| T1–Fagerström | 0.039 | **0.668<sup>\*\*\*</sup>** | 0.027 | **–0.105<sup>\*</sup>** | — |  |  |  |  |  |
| T1–Craving | **0.214<sup>\*\*\*</sup>** | **0.145<sup>\*\*</sup>** | **0.111<sup>\*</sup>** | **0.139<sup>\*\*</sup>** | **0.283<sup>\*\*\*</sup>** | — |  |  |  |  |
| T2–Tobacco-Pleasant<br>Preference | **0.548<sup>\*\*\*</sup>** | 0.048 | –0.068 | –0.126<sup>+</sup> | 0.047 | **0.224<sup>\*\*\*</sup>** | — |  |  |  |
| T2–Numbers of<br>Cigarettes | 0.073 | **0.880<sup>\*\*\*</sup>** | –0.042 | **–0.140<sup>\*</sup>** | **0.661<sup>\*\*\*</sup>** | **0.138<sup>\*</sup>** | 0.053 | — |  |  |
| T2–Quit Attempts | –0.054 | –0.032 | **0.813<sup>\*\*\*</sup>** | **0.289<sup>\*\*\*</sup>** | 0.048 | **0.161<sup>\*\*</sup>** | –0.052 | –0.075 | — |  |
| T2–Craving | **0.286<sup>\*\*\*</sup>** | 0.117<sup>+</sup> | **0.151<sup>\*</sup>** | 0.083 | **0.264<sup>\*\*\*</sup>** | **0.717<sup>\*\*\*</sup>** | **0.348<sup>\*\*\*</sup>** | **0.169<sup>\*\*</sup>** | 0.115<sup>+</sup> | — |

Note: Quit motivation and Fagerström test were assessed only at T1.
$p < .10$ , $^* p < .05$ , $^{**} p < .01$ , $^{***} p < .001$ . Significant p-values at $p < .05$ are highlighted in bold characters.

Since choice of tobacco>pleasant images positively correlated with craving, we divided the smokers into groups of high and low craving individuals using a median split. We then re-analyzed their behavior in the probabilistic choice test. At both T1 and T2, there was no group main effect for craving level (T1: F (1, 1864) = 3.196, P = 0.074; T2: (F (1, 1136) = 0.3147, P = 0.57)); the main effects of picture type (T1: F (3, 1864) = 114.6, P < 0.0001; T2: F (3, 1136) = 109.2, P < 0.0001) and the interactions between craving level and picture type (T1: F (3, 1864) = 8.487, P < 0.0001; T2: F (3, 1136) = 11.04, P < 0.0001) were significant. Post-hoc comparisons revealed that the interaction at T1 was driven by higher tobacco choice in the high cravers, an effect that was significant also at T2, where lower pleasant picture choice in this group was also discernable. Together, results suggest that experiencing craving increases the probability of selecting tobacco images.

## DISCUSSION

In this study, we adapted the PIC task ^31^ for measuring tobacco-seeking behavior (choice of viewing tobacco-related as compared to other salient and neutral images) in a relatively large sample of cigarette smokers. We found that the choice to view tobacco images was higher, and the choice to view pleasant images was lower, in smokers than nonsmokers. The tobacco as compared to pleasant image choice was stable when tested one month apart in the smokers (tested with an ICC), demonstrating a good test-retest reliability. It also correlated with self-reported levels of craving (both cross-sectionally and longitudinally), potentially pointing to a common underlying neurobiological mechanism, but not with other measures of TUD (including number of cigarettes smoked, quit attempts and motivation, and severity of dependence using the Fagerström questionnaire). In the smokers, high as compared to low cravers drove the enhanced drug and decreased pleasant picture choice. Given the strong evidence for the role of cues and craving in addiction and in relapse ^40–44^, which has been confirmed by recent meta-analyses ^6,10,45^, these results suggest that the PIC task is both a reliable (i.e., could be used in longitudinal studies) and valid measure of core processes involved in drug seeking. In particular, results suggest that the PIC task provides a stable indicator of tobacco-seeking behavior that is linked to incentive motivation for tobacco, rather than other TUD severity measures or cumulative nicotine exposure. Therefore, this task could serve as a valuable tool for investigating drug “wanting” and seeking behavior in the context of TUD.

Our results showing elevated drug>pleasant choice in smokers compared to non-smokers on this tobacco-adapted PIC task are consistent with prior versions of the task that were developed for use in cocaine use disorder ^31^ and validated for methamphetamine and opioid use disorders ^33,34^. In these prior studies this biased choice pattern was driven by current as compared to abstinent users (the former commonly reporting more craving and recent drug use), correlating with concurrent drug use outside the lab and predictive of actual drug use six months into the future. Interestingly and as expected, in the current study this drug>pleasant choice correlated with craving, where high cravers (as compared to low cravers) drove the observed choice bias. In contrast to previous studies that used tasks requiring choices between two explicit options (drug or alternative reward), where the selection of tobacco images was correlated not only with the intensity of craving but also with measures of drug use dependence (notably, the reported number of cigarettes smoked per day)^22,25,26^, here we did not observe significant correlations with our other measures of severity of TUD. The difference between the PIC task and other simpler, binary-choice, tasks used in previous studies might be attributed to considerable variations in methods and participants’ clinical profiles encompassing differences in the recruitment procedures (such as on-campus or treatment-seeking patients vs. online recruitment), the environment where the tests were conducted (laboratory vs. home settings) and the control over time since subjects smoked the last cigarette (ranging from several hours of abstinence to no control). Alternatively, it is possible that the PIC test measures some processes that, while overlapping, are somewhat distinct from those evaluated by the other choice procedures. Taken together, our results suggest that drug biased choice in the PIC task in people with TUD is primarily related to craving (a declarative desire for drug), rather than overall dependence or duration/severity of use.

An important consideration and possible limitation of this study is that it was performed online, in a non-controlled environment. Participants with TUD were not formally diagnosed by a clinical rater, rather they were included based on self-reported cigarette use. There is also a potential for distraction and reduced engagement compared with a laboratory setting, although we used attention checks to minimize this possibility throughout our procedures. Nevertheless, even if effects were somewhat diluted due to the lack of experimental control compared to a clinical laboratory setting (which could have contributed to somewhat smaller effect sizes found here than we have observed in prior studies with the PIC task), this potential limitation is balanced by the notable strength of studying a large, representative sample where even smaller effects could have large impact, by informing on relevant psychological and behavioral correlates of TUD in the community. Second, the current study was conducted in current smokers not seeking to quit tobacco use, and thus the results may not generalize to treatment-seekers. Another technical consideration stemming from this online sampling is that we could not standardize smoking abstinence at the time of the test. Participants could have smoked immediately before (possibly during) or several hours before taking the tests, precluding our ability to account for the pharmacological effects of nicotine or withdrawal, shown to influence craving ^46,47^. Another possible limitation is that we selected participants who were 35 years or older and younger populations remain to be studied. In addition, smokers and non-smokers were tested at different times and the impact of the pictures included remains to be studied in larger and simultaneously studied cohorts. Finally, craving has been described at both the trait (some individuals experience more intense craving than others) and the state (craving fluctuates over time and can be modulated by context and cues) levels ^48^, yet showing strong correlations^49^ suggestive of their interaction. Here we measured craving using a questionnaire aimed at evaluating state craving ^35^, yet its stability across both time points suggest its trait effects. Future studies are needed to determine whether the PIC task is sensitive to manipulations of state craving (for example, stress-induced or withdrawal-induced craving) and assess the relative contribution of both trait and state craving on this task performance.

The key finding of this study is that the PIC task provides an objective, reliable and valid proxy measure for evaluating the impact of tobacco cues on the choice of viewing cigarette-related images among smokers. The biased choice on the PIC task was closely associated with the craving and desire to smoke, highlighting its relevance to understanding smoking behavior. Importantly, this pattern of a drug choice bias at the expense of alternative reinforcers is consistent with the Impaired Response Inhibition and Salience Attribution (I-RISA) model of addiction ^50^ and other influential theories in drug addiction including incentive motivation ^51^, aberrant learning of drug-related stimuli ^11,12^ and drug-induced allostasis ^52^, which are based on a rich animal literature. Our current results using behavioral drug-seeking, analogous to non-human self-administration models of addiction ^53^, allow a translational bridge that is not dependent on intact processes of self-awareness/insight or interoception (needed for accurate self-reports) ^54^. This behavioral tool for assessing tobacco image seeking could also be useful in the classification, diagnosis, and prognosis of human TUD. Specifically, future work can determine whether the PIC task can serve as a useful predictor of long-term clinical outcomes (e.g., maintaining abstinence) in treatment-seeking people with TUD.

## Data Availability

Data and statistical analyses are freely accessible via the Open Science Framework link: https://osf.io/n23rx/

