## Supplementary Figures for "Tobacco Images Choice and Its Association with Craving and Dependence in Cigarette Smokers"

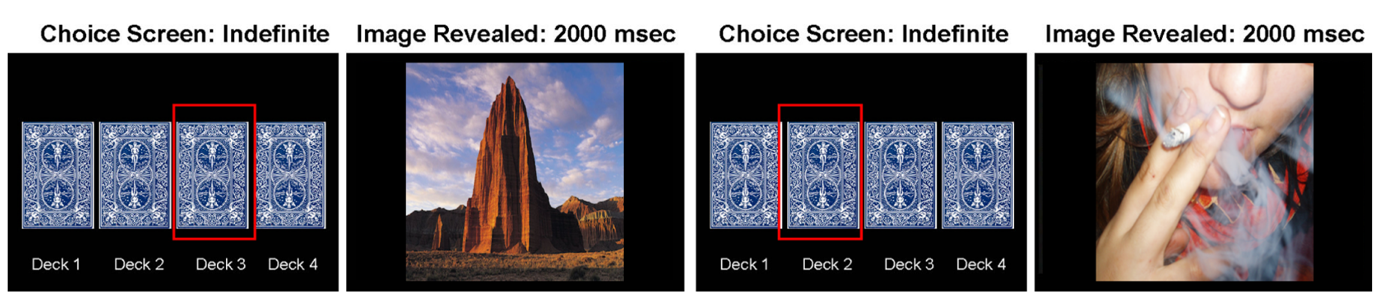

Fig. S1. Schematic representation of the adapted Probabilistic Image Choice (PIC) Task. Participants were instructed to select cards from 4 decks each containing a majority of cards of one of four categories: tobacco-related, pleasant, unpleasant, and neutral. Upon selection, the corresponding image was shown for 2000 ms. Participants were invited to select the most “appealing” images. A session consisted of 4 consecutive runs. If one deck was selected 8 times, the run ended and the position of decks was semirandomly changed.

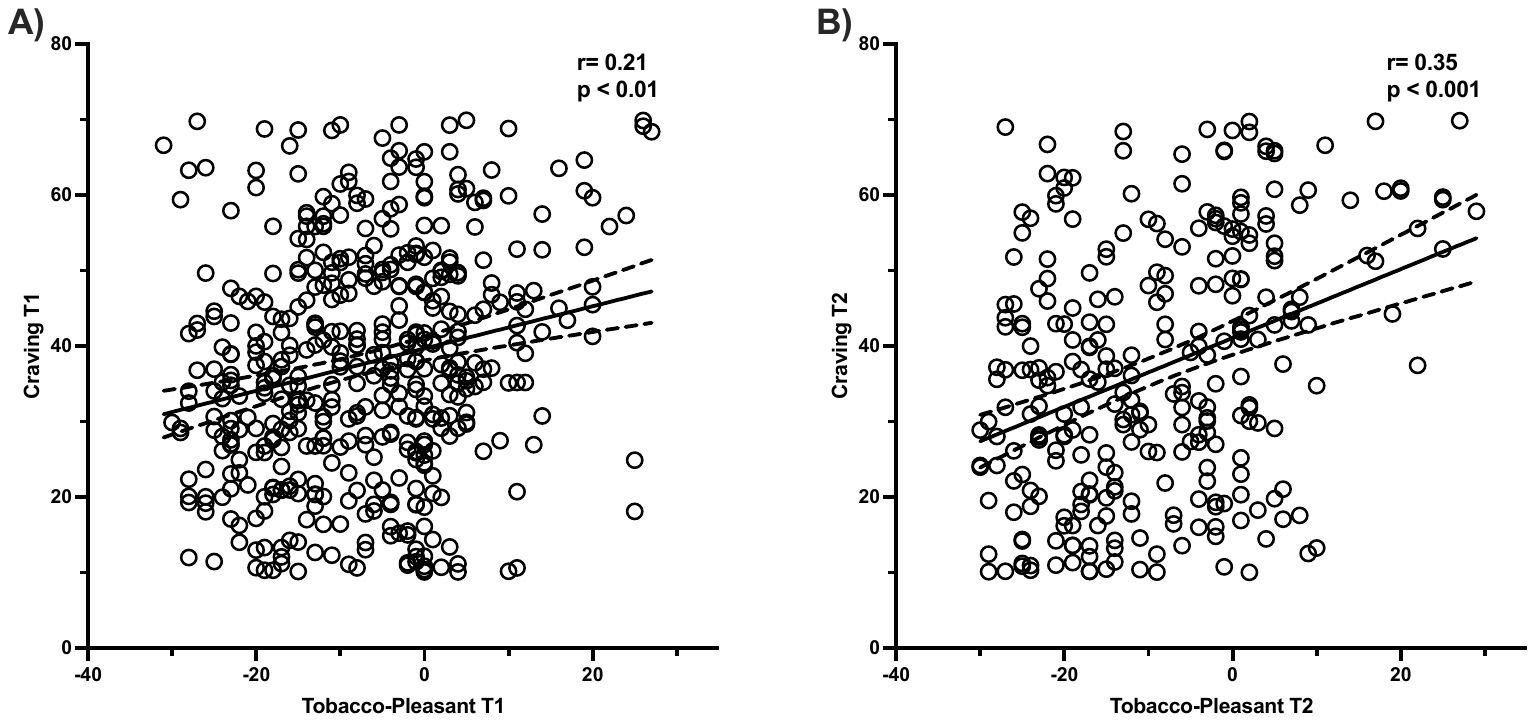

**Fig. S2. Correlation between preference for Tobacco images in the PIC task and self-reported craving at T1 (A) and T2 (B).** Correlation between the difference between the selection of tobacco images – pleasant images in the PIC task and craving scores measured at one-month interval. N = 286. Dotted lines show 95% CI.
